# Alterations of auditory sensory gating in mice with noise-induced tinnitus treated with nicotine and cannabis extract

**DOI:** 10.1101/2022.06.18.496668

**Authors:** Barbara Ciralli, Thawann Malfatti, Thiago Z. Lima, Sérgio Ruschi B. Silva, Christopher R. Cederroth, Katarina E. Leao

## Abstract

Tinnitus is a phantom sound perception affecting both auditory and limbic structures. The mechanisms of tinnitus remain unclear and it is debatable whether tinnitus alters attention to sound and the ability to inhibit repetitive sounds, a phenomenon also known as auditory gating. Here we investigate if noise exposure interferes with auditory gating and whether natural extracts of cannabis or nicotine could improve auditory pre-attentional processing in noise-exposed mice. We used 22 male C57BL/6J mice divided into noise-exposed (exposed to a 9-11 kHz narrow band noise for 1 hour) and sham (no sound during noise exposure) groups. Hearing thresholds were measured using auditory brainstem responses, and tinnitus-like behavior was assessed using Gap prepulse inhibition of acoustic startle. After noise exposure, mice were implanted with multi-electrodes in the dorsal hippocampus to assess auditory event-related potentials in response to paired clicks. The results showed that mice with tinnitus-like behavior displayed auditory gating of repetitive clicks, but with larger amplitudes and longer latencies of the N40 component of the aERP waveform. The combination of cannabis extract and nicotine improved auditory gating ratio in noise-exposed mice without permanent hearing threshold shifts. Lastly, the longer latency of the N40 component appears due to an increased sensitivity to cannabis extract in noise-exposed mice compared to sham mice. The study suggests that the altered central plasticity in tinnitus is more sensitive to the combined actions on the cholinergic and the endocannabinoid systems. Overall, the findings contribute to a better understanding of pharmacological modulation of auditory sensory gating.

## Introduction

Subjective tinnitus is a phantom sound sensation without an external source that is related to comorbidities such as anxiety and depression [1] and decreased quality of life [2]. Tinnitus affects around 15% of the world population [3] and so far cognitive behavioral therapy is the only evidence-based recommended treatment [4]. A relationship between tinnitus and decreased understanding of speech-in-noise has been reported [5] but it remains unclear whether chronic tinnitus directly interferes with speech-in-noise processing [6], or whether this is a result of attentional problems that have been difficult to assess in tinnitus subjects [5]. The limbic system is implicated in the manifestation and development of chronic tinnitus [7], and positron emission tomography (PET) and functional magnetic resonance imaging (fMRI) studies have shown greater activation of the auditory cortex, as well as non-auditory areas (frontal areas, limbic system and cerebellum) in tinnitus patients compared to controls [8]. Animal models of tinnitus point to neuronal alterations in the dorsal cochlear nucleus [9], affecting upstream auditory nuclei, with previous evidence of altered activity of the auditory cortex [10]. The auditory cortex has been shown to have significantly reduced functional connectivity with limbic structures (such as the hippocampus and amygdala) when comparing regional fMRI low-frequency activity fluctuations in a mouse model of noise-induced tinnitus [11]. Still, the involvement of limbic structures, such as the hippocampus, in noise-induced tinnitus remains poorly investigated.

Auditory information reaches the hippocampus through two distinct pathways: the lemniscal and non-lemniscal pathways, which converge in the entorhinal cortex before reaching the hippocampus [12]. Processing of auditory input in the hippocampus can be measured by auditory event-related potentials (aERP) for sensory gating, which is defined as a reduction in aERP to a repeated identical stimulus. Mouse aERP recordings are commonly performed on the CA1 and CA3 hippocampal regions [13, 14, 15]. Notably, the CA1 region maintains direct connections with the primary auditory cortex and auditory association areas [16]. This unique connectivity establishes the hippocampus as an important interface between the auditory and limbic systems, potentially impacted in neurological conditions such as tinnitus.

Auditory sensory gating can be assessed with paired-click stimuli (0.5 s apart) where the aERP magnitude in response to the second click generates a smaller amplitude compared to the first. In humans, aERPs are measured using electroencephalogram (EEG), while in mice aERPs are often recorded using intra-hippocampal chronically implanted electrodes [17, 13]. An incomplete suppression of the second click represents abnormal sensory processing, and poor “gating” of paired auditory stimuli [18]. A decrease in sensory gating measured by cortical aERPs in response to paired tones has been shown to be correlated with tinnitus severity in young adults [19], whereas an increased latency in aERP was found in tinnitus patients [20]. Still, the neuronal correlates of aERPs are poorly understood and animal models of noise-induced tinnitus measuring auditory gating are largely lacking even though the aERP waveform of rodents, described as positive (P) or negative (N) peaks, with approximate latency in milliseconds, P20, N40 and P80 [17] or P1, N1 and P2, are analogous to the human waveforms (P50, N100 and P200).

Pharmacologically it has been shown that certain nicotinic acetylcholine receptors take part in augmenting aERPs [17, 13]. Furthermore, it was shown that smoking cigarettes containing different doses of cannabis led to a reduction in the amplitude of event-related potentials. Additionally, subjects experienced an acutely diminished attention and stimulus processing after smoking cannabis [21]. On the contrary, a combined activation of the cholinergic and the endocannabinoid system has shown to improve auditory deviant detection and mismatch negativity aERPs in human subjects, but not when each drug was delivered alone [22]. This indicates interactions between the two systems, however, the impact of nicotine and/or cannabis, on aERPs in animal models of tinnitus, has to our knowledge not yet been studied. Here, we first hypothesized that noise-induced tinnitus interferes with auditory gating, and next that nicotine or natural extracts of cannabis could improve auditory pre-attentional processing in noise-induced tinnitus. To test this, we used a mouse model of noise-induced tinnitus without hearing impairment and measured aERPs in the dorsal hippocampus in response to paired clicks.

## Methods

### Animals

All protocols were approved by and followed the guidelines of the ethical committee of the Federal University of Rio Grande do Norte, Brazil (Comitê de Ética no Uso de Animais - CEUA; protocol no.094.018/2018). C57BL/6J male mice (1 month old at the beginning of the experimental timeline) originated from an in house-breeding colony. Here we used a total of 29 mice, where 7 were excluded in the Gap-prepulse inhibition of acoustic startle (GPIAS) test initial screening due to poor GPIAS (see exclusion criteria at the GPIAS section), leading to a total of 22 mice reported in all experimental procedures. Before the beginning of experiments, the animals were randomly assigned using python scripts (see section 2.11) to the Sham (n = 11) or Noise-exposed (n = 11) group. From those, 3 animals were excluded from aERP recordings due to low signal-to-noise ratio and 2 animals died after surgery (remaining 10 Sham and 7 Noise-exposed). Animals were housed on a 12/12h day/night cycle (onset/offset at 6h/18h) at 23ºC to maintain normal circadian rhythm and had free access to water and food pellets based on corn, wheat and soy (Nuvilab, Quimtia, Brazil;: #100110007, Batch: 0030112110). All experiments were performed during the day cycle, ranging from 7h to 15h. Animals (2-4 per cage) were housed in IVC cages, and paper and a polypropylene tube was added as enrichment. Once implanted, animals were single-housed until the end of the experiment. Mice were tunnel handled for the experiments as it has been shown to impact stress during experimental procedures, while tail-handling was used for routine husbandry procedures.

### Sound calibration

The sound equipment used for Auditory brainstem responses (ABRs), noise exposure, GPIAS and aERPs was calibrated in their respective arenas, all inside a sound-shielded room with background noise of 35 decibel sound pressure level (dBSPL). We used an ultrasonic microphone (4939-A-011, Brüel and Kjær) to record each of the stimuli used at 300 voltage steps logarithmically spaced in the 0-1V range, allowing to play all needed stimuli at the voltage necessary to achieve the needed intensity in dBSPL.

### Auditory brainstem responses

The ABRs of mice was tested both before and after the noise exposure protocol. Mice were anesthetized with an intraperitoneal injection (10 *μ*l/gr) of a mixture of ketamine/xylazine (90/6 mg/kg) plus atropine (0.1 mg/kg) and placed in a stereotaxic apparatus on top of a thermal pad with a heater controller (Supertech Biological Temperature Controller, TMP-5b) set to 37°C and ear bars holding in front of and slightly above the ears, on the temporal bone, to not block the ear canals. The head of the animal was positioned 11 cm in front of a speaker (Super tweeter ST400 trio, Selenium Pro). To record the ABR signal, two chlorinated electrodes were used, one recording electrode and one reference (impedance 1 kΩ) placed subdermally into small incisions in the skin covering the bregma region (reference) and lambda region (recording). Sound stimulus consisted of narrow-band uniform white noise pulses with length of 3 ms each, presented at 10 Hz for 529 repetitions at each frequency and intensity tested. The frequency bands tested were: 8-10 kHz, 9-11 kHz, 10-12 kHz, 12-14 kHz and 14-16 kHz. Pulses were presented at 80 dBSPL in decreasing steps of 5 dBSPL to the final intensity 45 dBSPL as previously described [23]. The experimenter was blinded to the animal group during the ABR recordings.

### Gap prepulse inhibition of acoustic startle

The GPIAS test [24] is known to reliably measure tinnitus-like behavior on rodents such as rats, mice and guinea-pigs [25, 26, 27, 28], and was used here to infer tinnitus in noise-exposed mice. GPIAS evaluates the degree of inhibition of the auditory startle reflex by a short preceding silent gap embedded in a carrier background noise. Before the first recording session, the animals were habituated to the experimenter and experimental setup for 3 consecutive days. Then, mice were acclimatized during the next 3 consecutive days by running the entire GPIAS session with all frequencies and trials but without the startle pulse. Animals were allowed 5 minutes inside the recording chamber before each recording session. Mice were then screened 3 days before the noise exposure for their ability to detect the gap. Animals were then tested again 3 days after noise exposure or sham procedures (no noise), as previously described [23]. Animals were placed in custom-made acrylic cylinders perforated at regular intervals. The cylinders were placed in a sound-shielded custom-made cabinet (44 × 33 x 24 cm) with low-intensity LED lights in a sound-shielded room with ≈35 dBSPL (Z-weighted) of background noise. A single loudspeaker (Super tweeter ST400 trio, Selenium Pro, freq. response 4-18 kHz) was placed horizontally 4.5 cm in front of the cylinder, and startle responses were recorded using a digital accelerometer (MMA8452Q, NXP Semiconductors, Netherlands) mounted to the base plate of the cylinder and connected to an Arduino Uno microcontroller, and a data acquisition cart (Open-ephys board) analog input. Sound stimuli consisted of 60 dBSPL narrow-band filtered white noise (carrier noise); 40 ms of a silent gap (GapStartle trials); 100 ms of interstimulus interval carrier noise; and 50 ms of the same noise at 105 dBSPL (startle pulse), with 0.2ms of rise and fall time. The duration of the carrier noise between each trial (inter-trial interval) was pseudo-randomized between 12-22 s. Test frequencies between 8-10, 9-11, 10-12, 12-14, 14-16 and 8-18 kHz were generated using a butterworth bandpass filter of 3rd order. The full session consisted of a total of 18 trials per frequency band tested (9 Startle and 9 GapStartle trials per frequency, pseudo-randomly played). It was previously shown that mice can suppress at least 30% of the startle response when the loud pulse is preceded by a silent gap in background noise [29], therefore we retested frequencies to which an animal did not suppress the startle by at least 30% in a second session the next day. Animals that still failed to suppress the startle following the silent gap in at least two frequencies in the initial GPIAS screening were excluded from further experiments. The experimenter was blinded to the animal group during the GPIAS recordings. Since we only assessed mice three days after noise exposure, while others suggest that chronic tinnitus arises after seven weeks in C57Bl6 mice [30], we infer our GPIAS relate to acute tinnitus.

### Noise exposure

Mice were anesthetized with an intraperitoneal administration of ketamine/xylazine (90/6 mg/kg), placed inside an acrylic cylinder (4 × 8 cm) facing a speaker (4 cm distance) inside a sound-shielded cabinet (44 × 33 x 24 cm) and exposed to a narrow-band white noise filtered (butterworth, -47.69dBSPL/Octave) from 9-11 kHz, at an intensity of 90 dBSPL for 1h. This protocol was previously shown to trigger a tinnitus-phenotype in C57BL/6 mice that could be decreased by chemogenetically modulating the firing rate of CaMKII*α*+ DCN units [23]. Next, mice remained in the cylinder inside the sound shielded chamber for 2 hours, due to the fact that sound-enrichment post loud noise exposure may prevent tinnitus induction [31]. Sham animals were treated equally, but without any sound stimulation. We used 11 noise-exposed and 11 sham animals. The animals were then returned to their home cages.

### Electrode array assembly

Tungsten insulated wires of 35 *μ*m diameter (impedance 100-400 kΩ, California Wires Company) were used to manufacture 2 × 8 arrays of 16 tungsten wire electrodes. The wires were assembled to a 16-channel custom made printed circuit board and fitted with an Omnetics connector (NPD-18-VV-GS). Electrode wires were spaced by 200 *μ*m with increasing length distributed diagonally in order to record from different hippocampal layers, such that, after implantation, the shortest wire were at dorsoventral (DV) depth of -1.50 mm and the longest at DV -1.96 mm. The electrodes were dipped in fluorescent dye (1,1’-dioctadecyl-3,3,3’,3’-tetramethylindocarbocyanine perchlorate; DiI, Invitrogen) for 10 min (for post hoc electrode position) before implanted into the right hemisphere hippocampus.

### Electrode array implantation

22 animals were used for the electrodes implantation surgery. In detail, mice were anesthetized using a mixture of ketamine/xylazine (90/6 mg/kg) and placed in a stereotaxic frame on top of a heat pad (37°C). Dexpanthenol was applied to cover the eyes to prevent ocular dryness. When necessary, a bolus of ketamine (45 mg/kg) was applied during surgery to maintain adequate anesthesia. Iodopovidone 10% was applied on the scalp to prevent infection, and 3% lidocaine hydrochloride was injected subdermally before an incision was made. In order to expose the cranial sutures, 3% hydrogen peroxide was applied over the skull. Four small craniotomies were done in a square at coordinates mediolateral (ML) 1 mm and anteroposterior (AP) -2.4 mm; ML: 1 mm and AP: -2.6 mm; ML: 2.45 mm and AP: -2.4 mm; ML: 2.45 mm and AP: -2.6 mm, to make a cranial window where the electrodes were slowly inserted at DV coordinate of -1.96 mm (for the longest shank). Four additional holes were drilled for the placement of anchoring screws, where the screw placed over the cerebellum served as reference. The electrode implant was fixed to the skull with polymethyl methacrylate moldable acrylic polymer around the anchor screws. After surgery, the animals were monitored until awake and then housed individually and allowed to recover for one week before recordings. For analgesia, ibuprofen 0.04 mg/ml was administered in the water bottle 2 days before and 3 days after the surgery. Subcutaneous Meloxicam 5 mg/kg was administered for 3 consecutive days after the surgery. 2 animals died shortly after the surgery, remaining 10 animals in the sham group and 7 in the noise-exposed group.

### Paired-click stimuli for auditory event related potentials

Mice were habituated during two days in the experimental setup and in the day of recording, anesthesia was briefly induced with isoflurane (5% for *<*1 min) to gently connect the implanted electrode array to a head-stage (Intan RHD 2132) connected to an acquisition board (OpenEphys v2.2 XEM6010-LX150) by a thin flexible wire. aERPs were recorded in freely moving animals placed in a low-light environment exposed to paired click stimulus, played by a speaker (Selenium Trio ST400) located 40 cm above the test area. All recordings were performed in standard polycarbonate cage bottom, which was placed inside a sound-shielded box (40 × 45 x 40 cm). The paired clicks consisted of white noise filtered at 5-15 kHz presented at 85 dBSPL, 10 ms of duration with 0.2 ms rise/fall ramp, and 0.5 s interstimulus interval. Stimulus pairs were separated by 2-8 s (pseudorandomly), and a total of 50 paired stimuli were presented. The session duration varied from 148 s to 442 s.

To investigate aERPs, average data from different animals, and also, compare responses from different experimental days and different pharmacological treatments, the appropriate hippocampal location for picking up aERP was identified. As local field potentials are related to cell density, and thereby the resistivity of the tissue, it is useful to record from the hippocampus with its distinct layered structure that shows phase-reversals of local field potentials [32]. Responses to paired clicks were recorded one week after surgery. The grand average of aERP (average of 50 clicks) for each channel was plotted and the changed signal polarity across hippocampal layers was identified, as the electrode array channels were distributed at different depths. To facilitate comparison of aERP between implanted animals we selected the first channel above phase reversal that showed a clear negative peak followed by a positive peak in the deeper channel. The visualization of the phase reversal channel was routinely added to analysis as channels sometimes shifted in the same animal, likely due to small movements in the electrode array when connecting/disconnecting mice to/from the headstage during different recording sessions. The experimenter was blind to the animal group during the aERP recordings.

### *Cannabis sativa* extract production and analysis

Δ^9^-tetrahydrocannabinol (THC) is the main psychoactive compound in cannabis and it is known to be partial agonist of cannabinoid receptor types 1 and 2 (CB1 and CB2) [33], while cannabinol (CBN) activates CB1 and CB2 receptors with more affinity over the latter and canabidiol (CBD) acts as a negative allosteric modulator of CB1 [33]. The Cannabis sativa extract was produced from an ethanolic extraction with the flowers previously dried and crushed. After leaving them in contact with the solvent for 5 min in an ultrasonic bath, filtration was performed and the process was repeated twice. Additionally, the solvent was evaporated and recovered, leaving only the cannabis extract in resin form. Decarboxylation of the acidic components, mainly tetrahydrocannabinolic acid into THC, was carried out by heating the material at 90°C until the conversion to the neutral forms had been completed. The cannabis extract was analyzed by high-performance liquid chromatography (HPLC). Analytical standards of THC (Cerilliant T-005), cannabinol (Cerilliant C-046) and CBD (Cerilliant C-045) were used in the calibration curve dilutions. An Agilent 1260 LC system (Agilent Technologies, Mississauga, ON, Canada) was used for the chromatographic analysis. A Poroshell 120 EC-C18 column (50 mm *×* 3.0 mm, 2.7 *μ*m, Agilent Technologies) was employed, with a mobile phase at a flow rate of 0.5 mL/min and temperature at 50°C (separation and detection). The compositions were (A) water and (B) methanol. 0.1% formic acid was added to both water and methanol. The total analysis time was 18 min with the following gradient: 0–10 min, 60–85%B; 10–11 min, 85–100%B; 11-12 min, 100%; 12-17 min, 100–60%; 17-18 min, 60% the temperature was maintained at 50°C (separation and detection). The injection volume was 5 *μ*L and the components were quantified based on peak areas at 230 nm. During the experiments we used a single dose of cannabis extract for each animal (100 mg/kg), containing 47.25 mg/kg of THC; 0.43 mg/kg of CBD and 1.17 mg/kg of CBN as analyzed by HPLC, and kindly donated by the Queiroz lab, Brain Institute, Federal University of Rio Grande do Norte, Brazil.

### Pharmacology

To activate the cholinergic system, and specifically brain nicotinic acetylcholine receptors, animals received a single intraperitoneal injection of nicotine (Sigma N3876) at 1.0 mg/kg [34] or saline (randomized order, 2 days in between session 1 and 2) 5 minutes before aERP recordings. In comparison to nicotine, which has a half-life of approximately 6-7 minutes in mouse plasma [35], THC, CBD, and CBN have longer half-lives. Specifically, THC has a half-life of approximately 110 minutes in mouse plasma [36], CBD has a half-life of 3.9 hours in mouse plasma [37], and CBN has a half-life of 32 hours in human plasma [38]. Here we administered a single dose of cannabis extract (100 mg/kg) and recorded responses after 30 minutes, similar to previously reported [39, 40, 41]. On the experimental day, the cannabis extract resin was diluted in corn oil to 10 mg/ml solution by mixing the extract and the oil and then sonicating for 5 min before injected intraperitoneally (at volume of 10 *μ*l/g body weight) 30 min prior to aERP recording sessions to reach max plasma concentration of THC [36]. After the third recording session, an additional dose of nicotine (1 mg/kg) was injected (to study potentially synergistic effects of cannabis extract + nicotine) and the animals were recorded 5 min later to observe how the interaction of the cholinergic and endocannabinoid system affects aERPs. After each aERP recording session, mice were unconnected from the headstage and returned to their home cage.

### Histology

To verify expected electrode positioning, animals were deeply anesthetized at the end of the experimental timeline with a mixture of ketamine/xylazine (180/12 mg/kg) and transcardiac perfused with cold phosphate buffered saline (PBS) followed by 4% paraformaldehyde (PFA). Brains were dissected and placed in 4% PFA for 48h. Next, brains were sliced using a free-floating vibratome (Leica VT1000S) at 75 *μ*m thickness, and cell nuclei were stained with 4’,6-diamidino-2-Phenylindole (DAPI, Sigma) to visualize cell layers and borders of the hippocampus. In addition to DiI-staining the electrodes, a current pulse of 500 *μ*A was routinely passed through the deepest electrode for 5 s at the end the last aERP recordings to cause a small lesion around the electrode tip to confirm electrode depth. Images were visualized using a Zeiss imager A2 fluorescence microscope with a N-Achroplan 5x objective.

### Data Analysis

Analysis of auditory brainstem responses was done as previously described [23] and consisted of averaging the 529 trials, filter the signal using a 3rd order butterworth bandpass filter from 600-1500 Hz, and slice the data 12 ms after the sound pulse onset. Thresholds were defined by automatically detecting the lowest intensity that can elicit a wave peak one standard deviation above the mean, and preceded by a peak in the previous intensity [23]. Effect of noise exposure and frequency of stimulus on ABR thresholds was evaluated using the Friedman Test, and pairwise comparisons were performed using the Wilcoxon signed-rank test. Effect of group was evaluated using the Kruskal-Wallis test, and pairwise comparisons were performed using the Mann-Whitney U test. Effect of group and frequency of stimulus on ABR threshold differences before and after exposure was evaluated using two-way analysis of variance (ANOVA). When multiple comparisons within the same dataset were performed, p values were Bonferroni-corrected accordingly.

For each frequency tested in GPIAS, Startle and Gap-Starle trials responses were separated and the signal was filtered with a Butterworth lowpass filter at 100 Hz. The absolute values of the accelerometer axes, from the accelerometer fitted below the cylinders enclosing the mice during the modified acoustic startle test, were averaged and sliced 400 ms around the startle pulse (200 ms before and 200 ms after). The root-mean-square (RMS) of the sliced signal before the Startle (baseline) was subtracted from the RMS after the startle response (for both Startle only and GapStartle sessions). The GPIAS index for each frequency was then calculated as

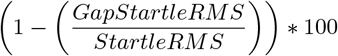

generating percentage of suppression of startle. For each animal, the most affected frequency was determined as the frequency with the greatest difference in GPIAS index before and after noise exposure. This was done as mice did not show decreased GPIAS at the same narrow-band frequency despite being subjected to the same noise exposure, indicating individual differences in possible tinnitus perception [26]. The definition of the most affected frequency followed the same procedure for both sham and noise-exposed animals. The effects of group (sham or noise-exposed), epoch (before or after exposure) and frequency of stimulus were tested using 3-way mixed models ANOVA. The effect of the group and epoch on the GPIAS index of the most affected frequency was evaluated using the Kruskal-Wallis and the Friedman test, respectively; and pairwise comparisons were done using the Mann-Whitney U and Wilcoxon signed-rank test, respectively.

Auditory event-related potentials in response to paired-clicks were filtered using a low pass filter at 60 Hz, sliced 0.2 s before and 1 s after the first sound click onset, and all 50 trials were averaged. To compare signals between different animals (n = 10 sham and n = 7 noise-exposed) and different treatments we always analyzed the channel above hippocampal phase reversal with a negative peak around 40 ms (N40) and a positive peak around 80 ms latency (P80). aERP components were quantified by peak amplitude (baseline-to-peak) after stimulus onset. The N40 was considered as the maximum negative deflection between 20 and 50 ms after the click stimulus, and P80 as the maximum positive deflection after the N40 peak. The baseline was determined by averaging all 50 trials and then averaging the 200 ms of prestimulus activity (before the first click). The latency of a component was defined as the time of occurrence of the peak after stimulus onset. The ratio in percentage of the first and second click amplitude (the suppression of the second click, e.g. sensory filtering) was calculated as

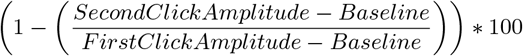

and error bars represent standard error of the mean (s.e.m) for all figures. A gating improvement was considered when the aERP peak amplitude suppression ratio(s) increased compared to sham (when comparing between groups) or to saline (when comparing between treatments). Effect of group, treatment and click on amplitude and latency of aERP components were evaluated using 3-way mixed-models ANOVA; effect of group and treatment in aERP components suppression, delay and N40-P80 width were evaluated using 2-way mixed-models ANOVA; and Student’s t-test was used for pairwise comparisons. Whenever the response failed to comply with normality, homoscedasticity and independence assumptions and parametric fitting was inadequate, the Kruskal-Wallis test was used to evaluate the effect of group, and the Mann-Whitney U test was used for pairwise comparisons; and the Friedman test was used to evaluate the effect of treatment and click, and Wilcoxon signed-rank test was used for pairwise comparisons. Statistical power for the tests ranged from 78.5 to 92.2%, and post-hoc multiple comparisons were adjusted by Bonferroni correction. Differences in occurrence of double-peak responses were evaluated using McNemar’s test.

## Results

In order to investigate whether noise-exposure can affect auditory gating we established an experimental timeline for experiments evaluating auditory perception using three different tests in mice exposed to a mild noise (90 dB-SPL, 9-11 kHz, 1h): ABRs, GPIAS and aERPs. Hearing thresholds of mice were assessed using ABRs 2 days before (baseline) and 2 days after sham or noise exposure (Figure 1A). ABRs showed field potentials with distinct peaks indicating neuronal activity at the auditory nerve, cochlear nuclei, superior olivary complex, and inferior colliculus [42] in response to sound clicks presented at different frequencies (Figure 1B-C). Similar to sham, noise exposure did not cause any change in ABR hearing thresholds at all frequencies tested when compared to baseline (Group, Kruskal-Wallis eff. size = 4.1e-05, p = 0.923; Epoch, Friedman eff. size = 0.058, p = 0.08; Frequency, Friedman eff. size = 0.007, p = 0.164; Figure 1D). When plotting threshold shifts, we confirmed that noise-exposed animals were impacted to a similar degree than sham mice (ANOVA; Group, F(1,21) = 0.047, p = 0.83; Frequency, F(4,84) = 0.2, p = 0.938; Group:Frequency, F(4,84) = 2.021, p = 0.09; Figure 1E). Unlike other models of tinnitus [43], we did not detect any effect of noise exposure in ABR Wave 1 amplitude (Epoch, Friedman test, eff. size = 0.037, p = 0.118; Group, Kruskal-Wallis test, eff. size = 0.0002, p = 0.821) or Wave 5 latency (Epoch, Friedman eff. size = 0.002, p = 0.55; Group, Kruskal-Wallis eff.size = 0.014, p = 0.073, Supplemental Figure S1). These findings confirm that the noise exposure did not cause any detectable change in hearing thresholds, and suggest a negligible impact on cochlear synaptopathy.

**Figure 1:**
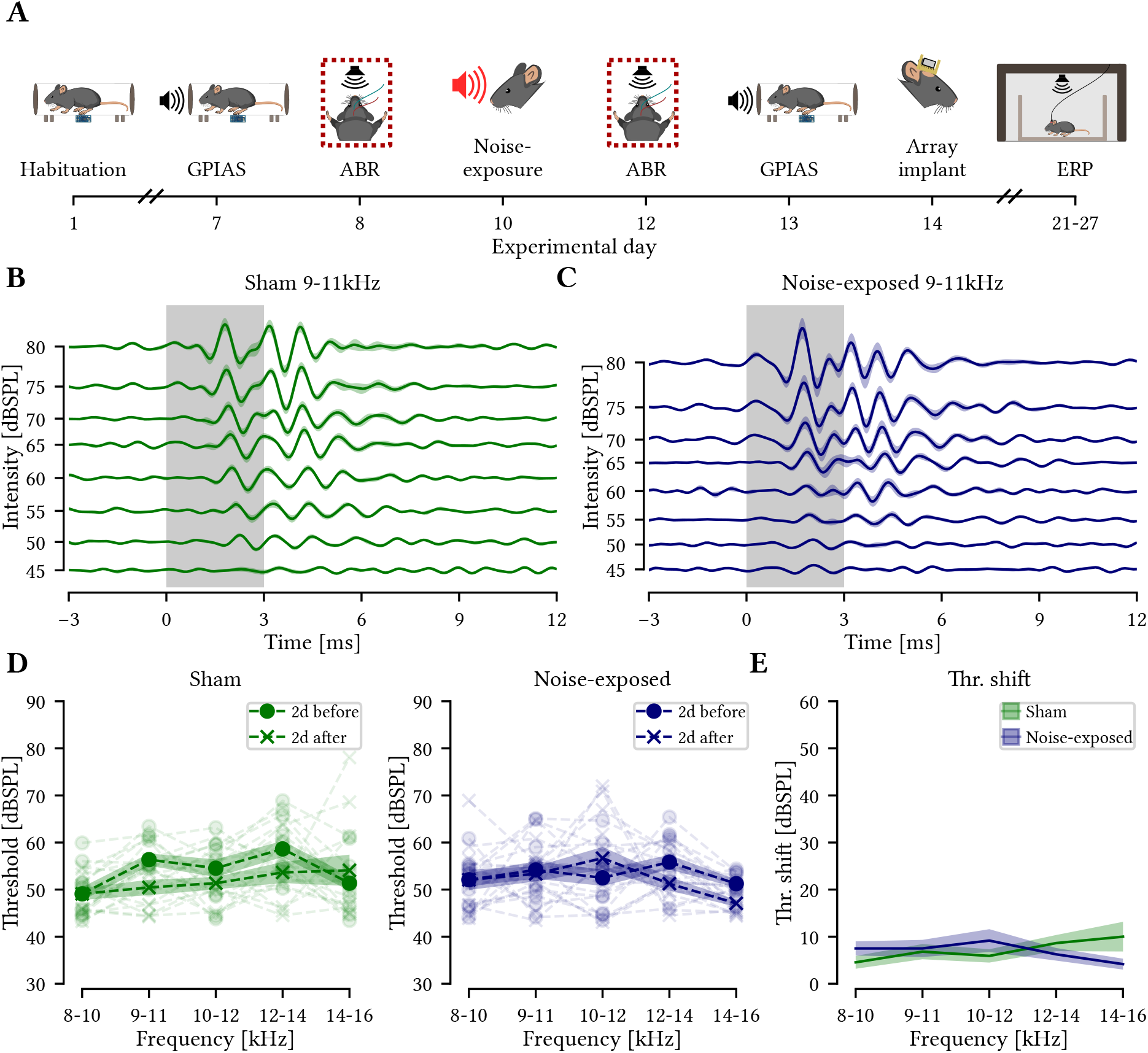
Noise exposure did not cause hearing threshold shift. A) Full experimental timeline highlighting time of ABR recordings (dotted rectangle). B-C) Mean auditory brainstem response (ABR) to 9-11kHz after noise-exposure for intensities 45-80 dBSPL for all 529 trials of all sham mice (B) and noise-exposed animals (C). Shaded traces show SEM, gray square indicates the sound pulse duration. D) Mean+SEM (line and shade) displaying auditory thresholds quantified for sham (n = 11, left) and noise-exposed (n = 11, right) animals two days before and two days after noise exposure, showing no significant difference at any frequency tested (Wilcoxon signed-rank test, p *>* 0.05 for all frequencies in both groups). E) Mean+SEM (line and shade) threshold shift for sham and noise-exposed mice showing no significant difference between groups at any frequency (Student’s t-test, p *>* 0.05 for all frequencies).

Three days before and 3 days after noise exposure mice were tested for GPIAS (Figure 2A-C). No effect of group (sham or noise-exposed), epoch (before or after noise exposure procedure) or frequency of stimulus was found in GPIAS when evaluating all frequencies from every animal (the closest to significance being the stimulus frequency factor; F(5,65) = 1.419, p = 0.229; Figure 2D-E) and no pairwise differences between any group, epoch or frequency, possibly due to each individual mouse may experience a different tinnitus pitch. We therefore evaluated the background frequency that interferes most with gap prepulse startle suppression for each individual mouse, which would correspond to the most likely tinnitus pitch of these animals (Figure 2F-G). Sham exposure had no effect on GPIAS (Friedman test; eff.size = 0.075; p = 0.365; Figure 2F, left), while in noise-exposed mice the noise exposure had a significant effect in GPIAS index (Friedman test; eff. size = 1.0; p = 1.8e-03), showing a decrease in startle suppression when comparing before and after noise exposure (Wilcoxon signed-rank test, p=9.8e-04; Figure 2F, right). Accordingly, the group (sham vs noise-exposed) had a significant effect on GPIAS measured after noise exposure (Kruskal-Wallis; eff. size = 0.663, p = 3.2e-04), with noise-exposed mice showing lower GPIAS suppression than sham mice (Mann-Whitney U; eff. size = 0.805; p = 4.0e-05); but not before noise exposure (Kruskal-Wallis; eff. size = 0.117, p = 0.066). GPIAS showed individual variability in the most affected frequency (Figure 2G), consistent with previous reports [26] and confirms that tinnitus interferes with the ability to suppress the startle response in noise-exposed animals.

**Figure 2:**
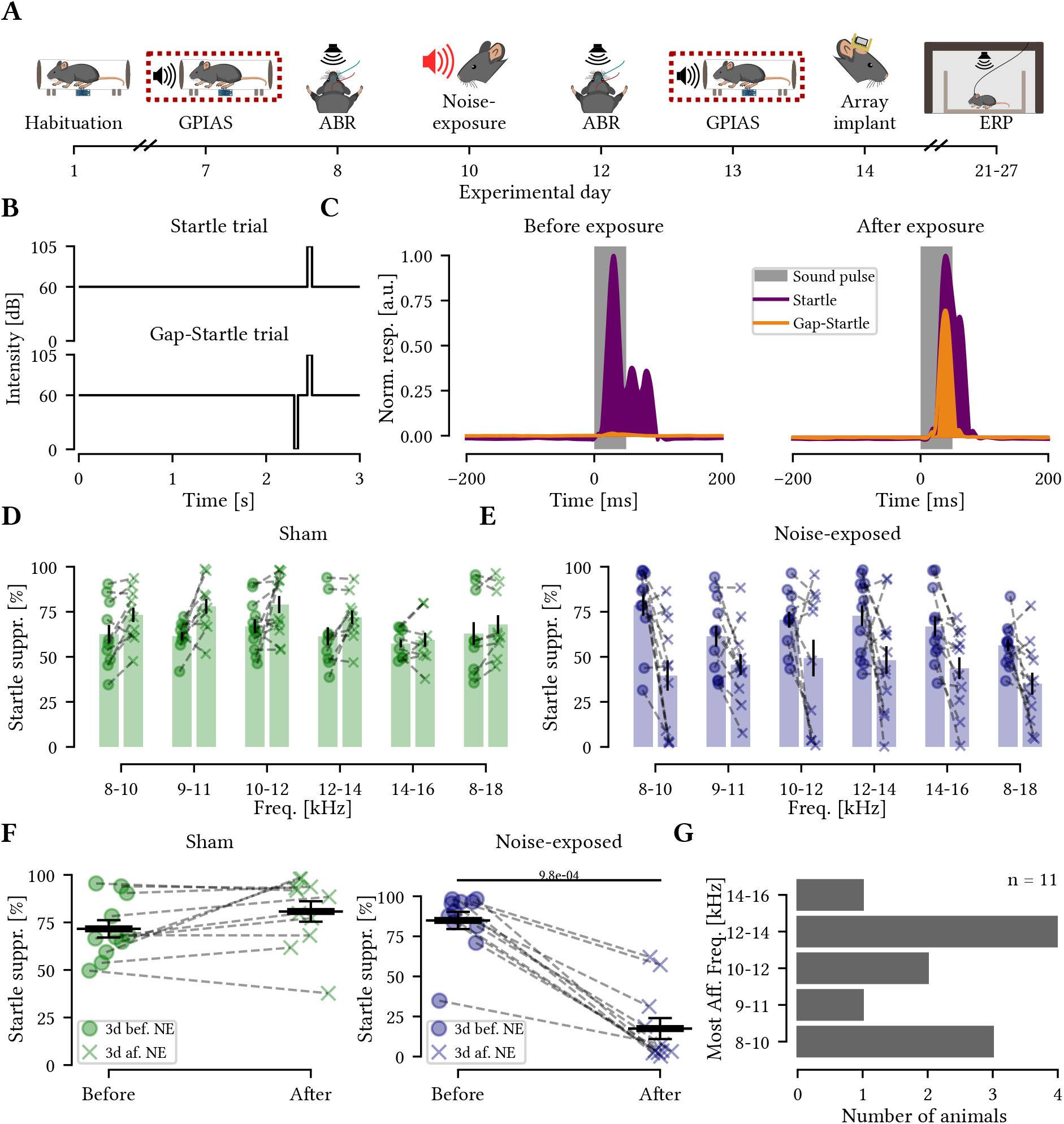
Noise-exposed animals showed decreased startle suppression. A) Timeline of experiments highlighting time point of the GPIAS tests. B) Schematic GPIAS protocol. C) Representative examples of startle suppression by the gap (left) and negative startle suppression (right) from the same animal 3 days before and 3 days after noise exposure, respectively. Filled traces represent an average of 9 trials of stimulus without gap (purple) and with gap (orange). Gray rectangle represents the 50ms startle stimulus. D-E) GPIAS index for all frequencies tested 3 days before (o) and 3 days after (x) noise exposure for sham (D) and noise-exposed (E) mice. F) The frequency with largest difference in startle suppression before and after noise-exposure was used for quantification of group GPIAS performance. Sham animals show no difference in GPIAS performance before and after noise exposure (left, n=11), while noise-exposed mice (right) show a significant decrease in startle suppression by the silent gap (Wilcoxon signed-rank test, n = 11, p = 9.8e-04). G) The frequency with largest difference in startle suppression before and after noise-exposure varied between individual noise-exposed mice.

After the ABR and GPIAS tests, electrodes were implanted in the dorsal hippocampus for the assessment of sensory gating (Figure 3A). As expected, auditory event-related potential recordings showed that the second click consistently generated a smaller aERP (Figure 3B) and the magnitude of peaks around 40ms and 80ms were quantified from baseline as the N40 and P80 peak, respectively, for both the first and second click in the phase-reversal channel (see Methods, Figure 3B-C). Next, to investigate the impact of noise-induced tinnitus on auditory gating (11 days after noise-exposure), freely exploring mice were individually subjected to randomized paired-click stimuli where both sham and noise-exposed mice presented characteristic aERP (Figure 3D). Two types of measurements were evaluated: the responses to sound clicks measured in the hippocampus (amplitude in *μ*V and latency in ms), which is a measurement of sound processing in the limbic system; and the ratio between the second and the first click responses (both amplitude and latency unitless), which measures the sensory gating.

**Figure 3:**
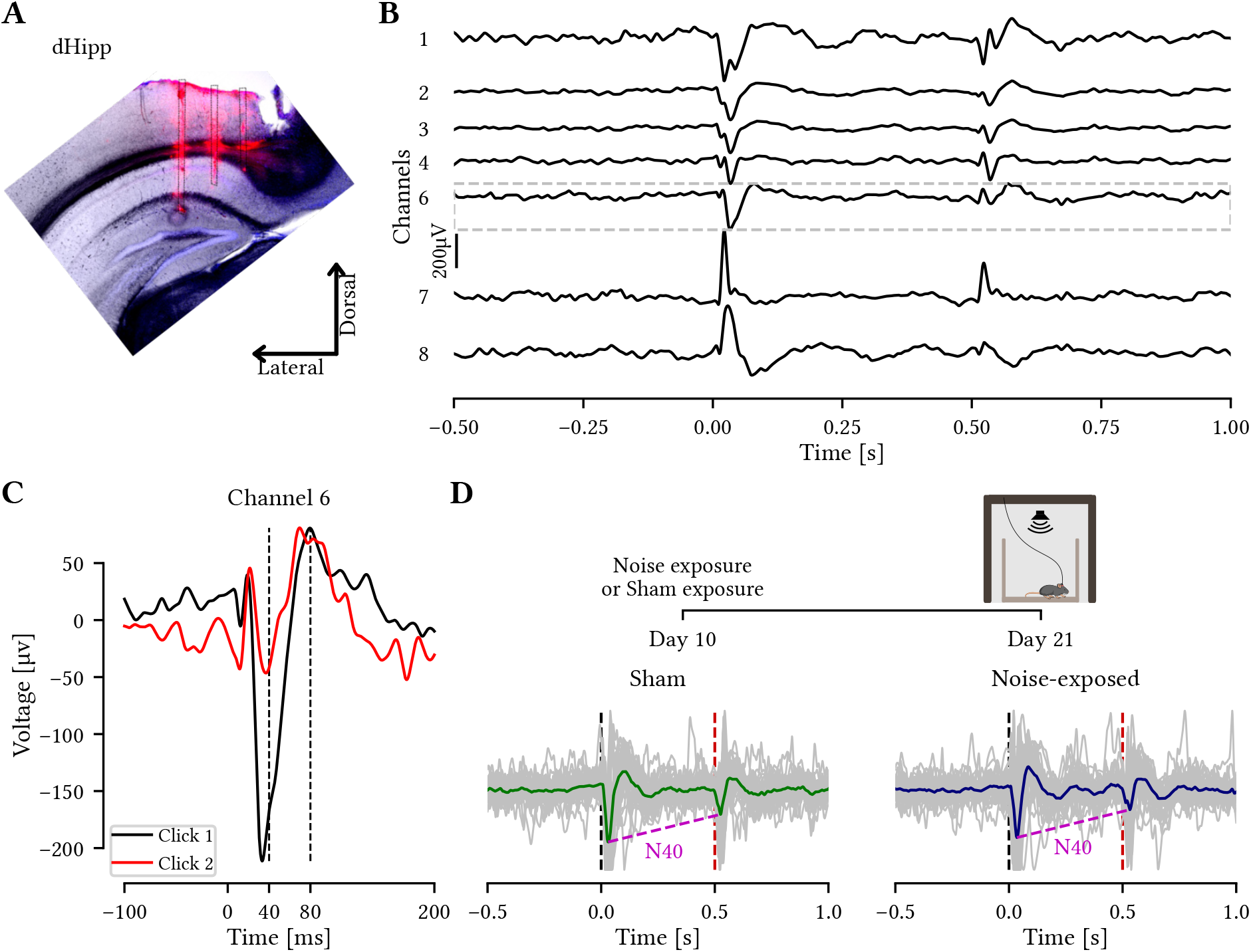
Auditory event-related potentials in sham and noise-exposed mice. A) Left, image of a mouse implanted with an electrode array. Right, coronal slice showing the dorsal hippocampus with electrode tracts stained with DiI in the CA1 region. B) Average aERPs in response to paired clicks from 8 channels at different depths from a recording session from a single animal. The channel above phase reversal (gray dotted box) was consistently used for aERP quantification. C) The reversal channel from ‘B’ at a greater magnification with click 1 (black) and 2 (red) responses superimposed. Dashed lines indicating positive and negative peaks at different characteristic latencies (N40 and P80 components). D) Top, simplified experimental timeline. Bottom, average traces of click responses in saline condition for sham (green, n = 10) and noise-exposed animals (blue, n = 7). Superimposed gray traces are the average response of 50 trials from each individual animal, dashed lines indicate the sound stimuli onset and amplitude difference of N40 peaks.

As attention is modulated by the cholinergic system [44] and also the endocannabinoid system [45], we tested the impact of two agonists to both systems (nicotine and cannabis extract, individually or in combination) in modulation of aERPs in our model of noise-induced tinnitus (Figure 4A). Animals were given a single injection of nicotine (1 mg/kg) or saline before aERP recordings on the fisrt two sessions. During the third session, the remaining two aERP recordings were conducted, with the initial recording taking place 30 minutes after the administration of cannabis extract (100 mg/kg). Subsequently, an additional dose of nicotine (1 mg/kg) was injected to investigate the potential synergistic effects of combining cannabis extract with nicotine. The average of the N40 response in sham-exposed animals showed the second click to be consistently smaller in amplitude compared to the first click (F(1,10) = 29.9, p = 2.7e-04; Supplemental Figure S2A, left). This significant attenuation on the second click was also observed for noise-exposed (F(1,10) = 11.2, p = 7e-03; Supplemental Figure S2A, right). The second click attenuation differed in strength depending on the pharmacological treatment between sham and noise-exposed mice (F(3,60) = 3.67, p = 1.7e-02; Supplemental Figure S2A). For noise-exposed animals the second click response was decreased compared to the first in nicotine (p = 1.6e-02) and cannabis extract + nicotine (p = 1.6e-02) treatment but not in saline (p = 0.237) or cannabis extract alone (p = 0.216 ; Supplemental Figure S2A, right), in contrast to sham animals. We thereby found a significant interaction between treatment and animal condition (sham or noise-exposed) on the N40 suppression ratio (F(3,60) = 3.5, p = 2e-02, Figure 4B). Looking specifically at sham mice, no significant difference was found in the N40 aERP ratio between treatments, while for noise-exposed animals, pairwise comparisons showed an increased N40 amplitude ratio after administration of cannabis extract + nicotine compared to cannabis extract alone (p = 1.9e-02), nicotine alone (p = 3.2e-02) and NaCl treatment (p = 1.9e-02, Figure 4B). There was also a significant difference in N40 ratio under cannabis extract + nicotine treatment between sham and noise-exposed mice (p = 1.0e-02; Figure 4B). We found a general effect of group in the N40 amplitude, where noise-exposed animals consistently showed a greater average when compared to sham-exposed mice (F(1,20) = 7.467; p = 6.3e-03; Figure 4C, Supplemental Table S1). Taken together, these results indicate that nicotine has a more pronounced effect on the filtering of repetitive stimuli in noise-exposed animals compared to sham animals, and that the combination of nicotine + cannabis extract strongly enhances the first 4C, Supplemental Table S1). Taken together, these results indicate that nicotine has a more pronounced effect on the filtering of repetitive stimuli in noise-exposed animals compared to sham animals, and that the combination of nicotine + cannabis extract strongly enhances the first and second click ratio in noise-exposed animals, an effect not seen in sham animals.

**Figure 4:**
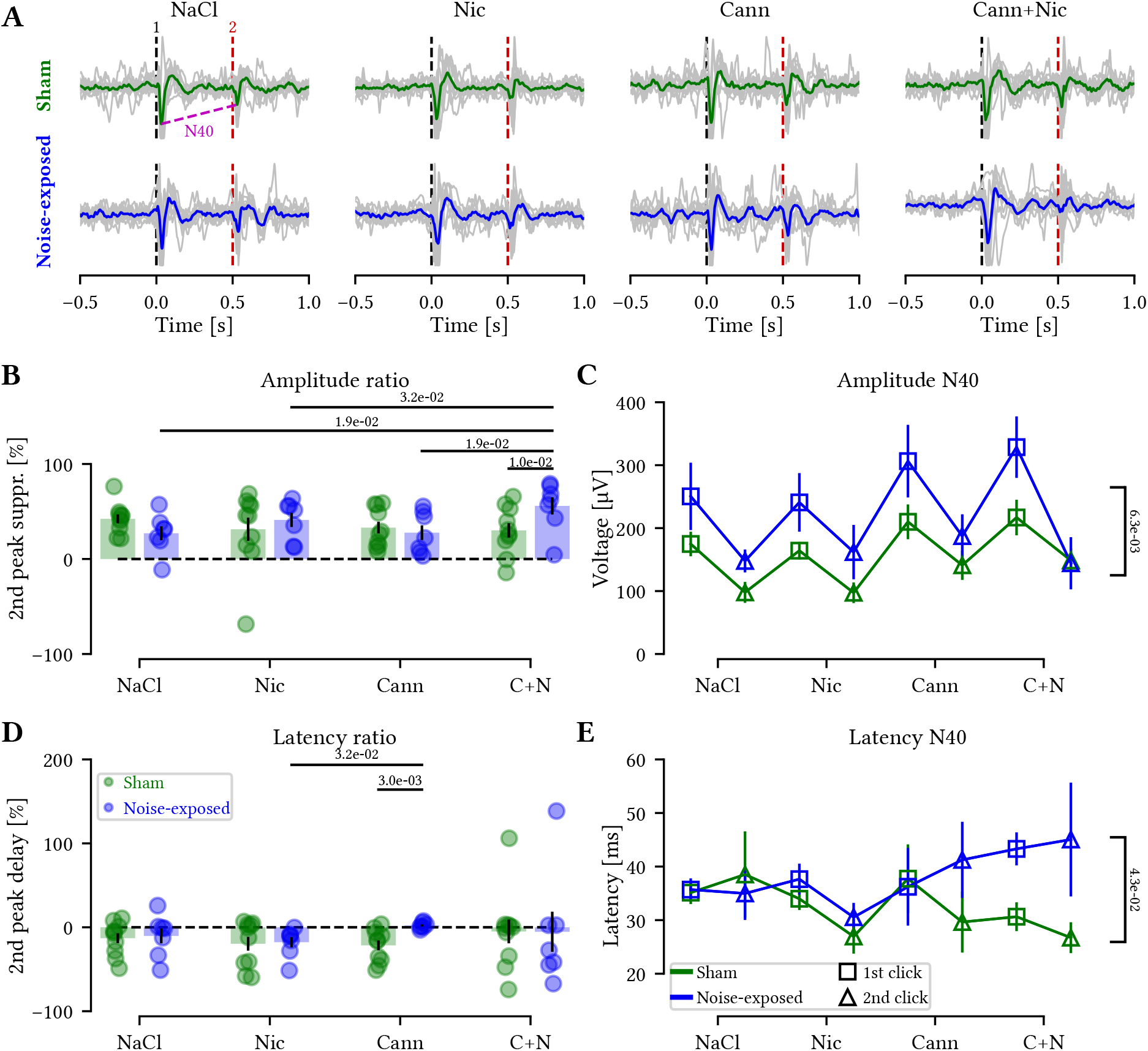
Noise-exposed mice have improved auditory gating under cannabis+nicotine treatment and showed overall larger and slower ERP responses. A) Auditory ERP recorded in awake mice in response to saline, nicotine, cannabis and cannabis+nicotine show characteristic suppression of the second click in both sham (top) and noise-exposed (bottom) animals. Gray trace shows the average aERP per animal while the green and blue traces show the group average for each treatment. B) Percentage of suppression of the second click of the N40 component (supplementary Figure S2) for sham (green) and noise-exposed (blue) mice, showing largest suppression of the second peak in noise-exposed mice following cannabis+nicotine administration (Student’s t-test). C) N40 amplitude is consistently increased for noise-exposed (n = 7) compared to sham animals (n = 10). D) Percentage of the second N40 peak delay for both groups at each treatment showed cannabis extract to increase delay in noise-exposed mice compared to sham (Mann-Whitney U test), as well as compared to nicotine treatment of noise-exposed mice (Wilcoxon signed-rank test). E) N40 latency is consistently increased for noise-exposed (n = 7) compared to sham animals (n = 10).

Examining latency of the N40 component showed no differences in pairwise comparisons between clicks after any particular treatment (p*>*0.05; Supplemental Figure S2B) although the distribution of latencies showed the second N40 latency to be consistently shorter compared to the first (p = 2.6e-03, Friedman test). Comparing the ratio of the first and second click latency revealed an increased response-delay in noise-exposed animals under cannabis treatment compared to sham animals in the same treatment (p = 3.0e-03) and compared to noise-exposed mice after nicotine administration (p = 3.2e-02; Figure 4D). This shows that cannabis delays the N40 latency compared to nicotine in noise-exposed animals but not in sham animals (Figure 4D). Overall, an effect of group on latency was found, where latency was consistently increased for noise-exposed mice (p = 4.3e-02, Kruskal-Wallis test; Figure 4E, Supplemental Table S2).

The P80 component of auditory aERP has been implicated in the NMDA dysfunction theory in schizophrenia, as ketamine can alter the P80 amplitude of mice [46]. The P80 component in response to the second click was consistently smaller compared to the response to the first stimulus (F(1,20) = 6.156, p = 2.2e-02). Also, the latency for the peak was reduced by the repetition of stimuli for both groups and all treatments (F(1,20) = 9.79, p = 5.2e-03). However, pairwise comparisons did not show any statistical differences for the P80 baseline to peak amplitude or latency (Figure 5A; Supplemental Figure S3) nor in ratios between the two clicks for the P80 amplitude (Figure 5B) and latency (Figure 5C). This indicates that the P80 component is not affected by noise-induced tinnitus.

**Figure 5:**
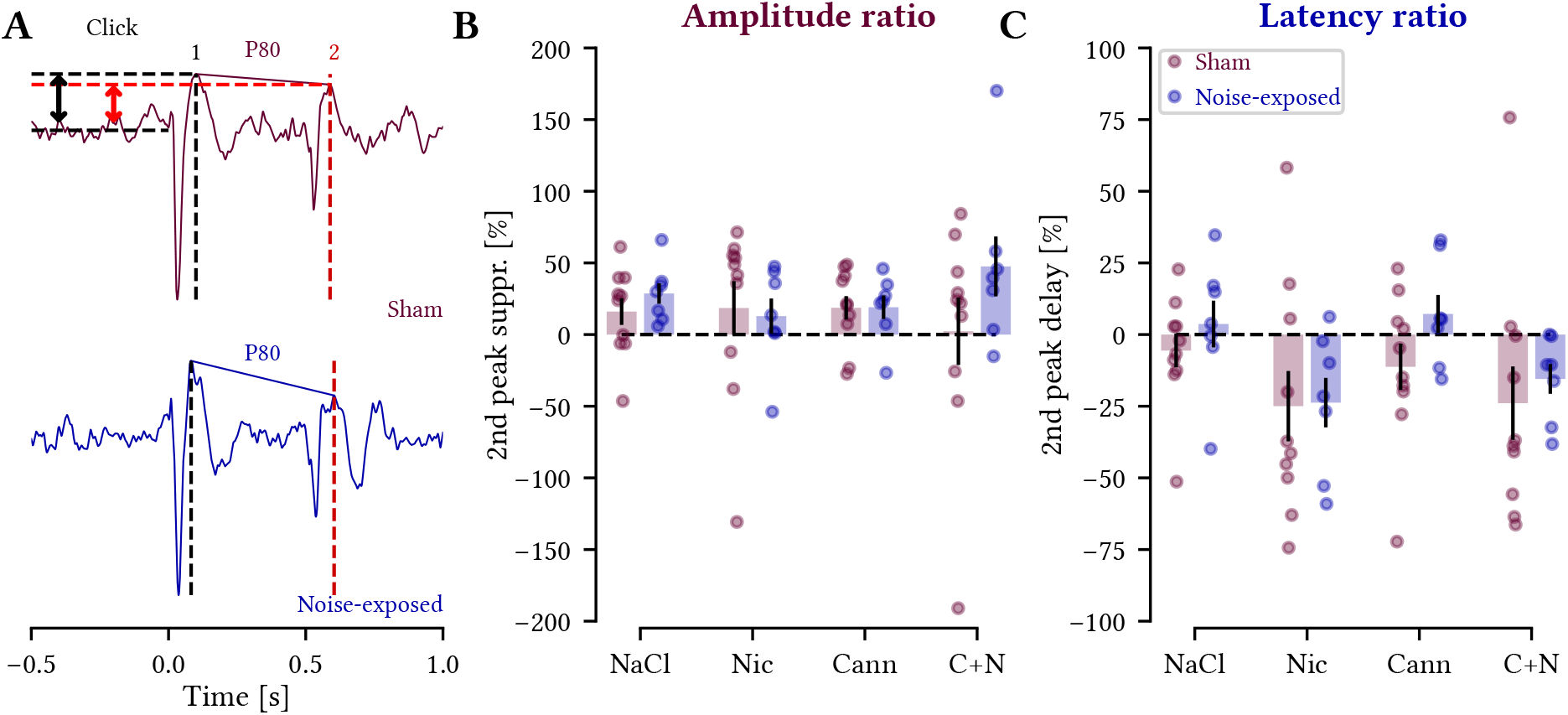
The P80 aERP amplitude and latency was not affected by noise-exposure or by nicotine and/or cannabis extract treatment. A) Representative trace highlighting the P80 component (vertical black and red dashed lines for first and second clicks, respectively). Arrows represent the calculated amplitude for each P80 response for the top trace. B) The percentage of second peak amplitude suppression showed no difference between sham and noise-exposed mice. C) Second P80 peak delay (ratio of the 1st and 2nd click responses latencies) for sham (purple) and noise-exposed (blue) animals showed no difference between groups or treatments. A negative ‘delay’ refers to a peak advancement. Wilcoxon signed-rank test, n = 10 sham and 7 noise-exposed mice, p *>* 0.05 for all comparisons.

As previous studies suggested that the improvement of sensory gating by pharmacological agents is mediated by an enhancement of the first click rather than by the suppression of the second click [17, 13], we separated the analysis of aERPs to focus on each click response (first; click 1 and repeated; click 2) by comparing the amplitude and latency of the N40 or P80 components between different treatments (Figure 6; Supplemental Figure S4). First, we found that sham animals increased the response to the first click after cannabis extract + nicotine treatment compared to just nicotine administration (p = 4e-03; Figure 6A, top left). Next, examining the repeated click 2 response, showed that the cannabis extract increased the N40 click 2 response amplitude compared to nicotine (p = 2.7e-02) and cannabis extract + nicotine also increased the N40 click 2 amplitude compared to nicotine alone (p = 6e-03; Figure 6A, top right). For the noise-exposed group, the combination of cannabis extract + nicotine increased click 1 amplitude compared to NaCl (p = 1.2e-02; Figure 6A, bottom left). There was no increase in click 1 response by nicotine, but still nicotine had an effect in the combination of cannabis extract since the combination of the two increased the response amplitude significantly compared to cannabis extract alone (p = 4.7e-02; Figure 6A, bottom left). The second click was unaltered by nicotine and/or cannabis extract for noise-exposed mice (Figure 6A, bottom right). Examining the latency of the N40 response to the first click showed no alteration by either treatment in the sham group (Figure 6B, top left). For the repeated click 2 latency, the sham group instead showed decreased latency in the presence of cannabis extract compared to NaCl treatment (p = 1.4e-02; Figure 6B, top right). For the noise-exposed group, cannabis extract + nicotine significantly delayed the click 1 N40 response compared to NaCl (p = 3.1e-02; Figure 6B, bottom left). Again, the latency of the second click N40 response was not affected by nicotine and/or cannabis extract in noise-exposed mice (Figure 6B, bottom right). Next, examining the P80 amplitude and latency in detail only showed one effect on the second click latency for noise-exposed mice where cannabis extract + nicotine marginally increased the latency of P80 click 2 response compared to nicotine alone (p = 4.9e-02; Supplemental Figure S4). All together we found the repeated second click N40 response to not be consistently modulated by treatment in noise-exposed mice, thereby agreeing with previous literature that pharmacological improvement of sensory gating affects the first click response for this set of animals [17, 13].

**Figure 6:**
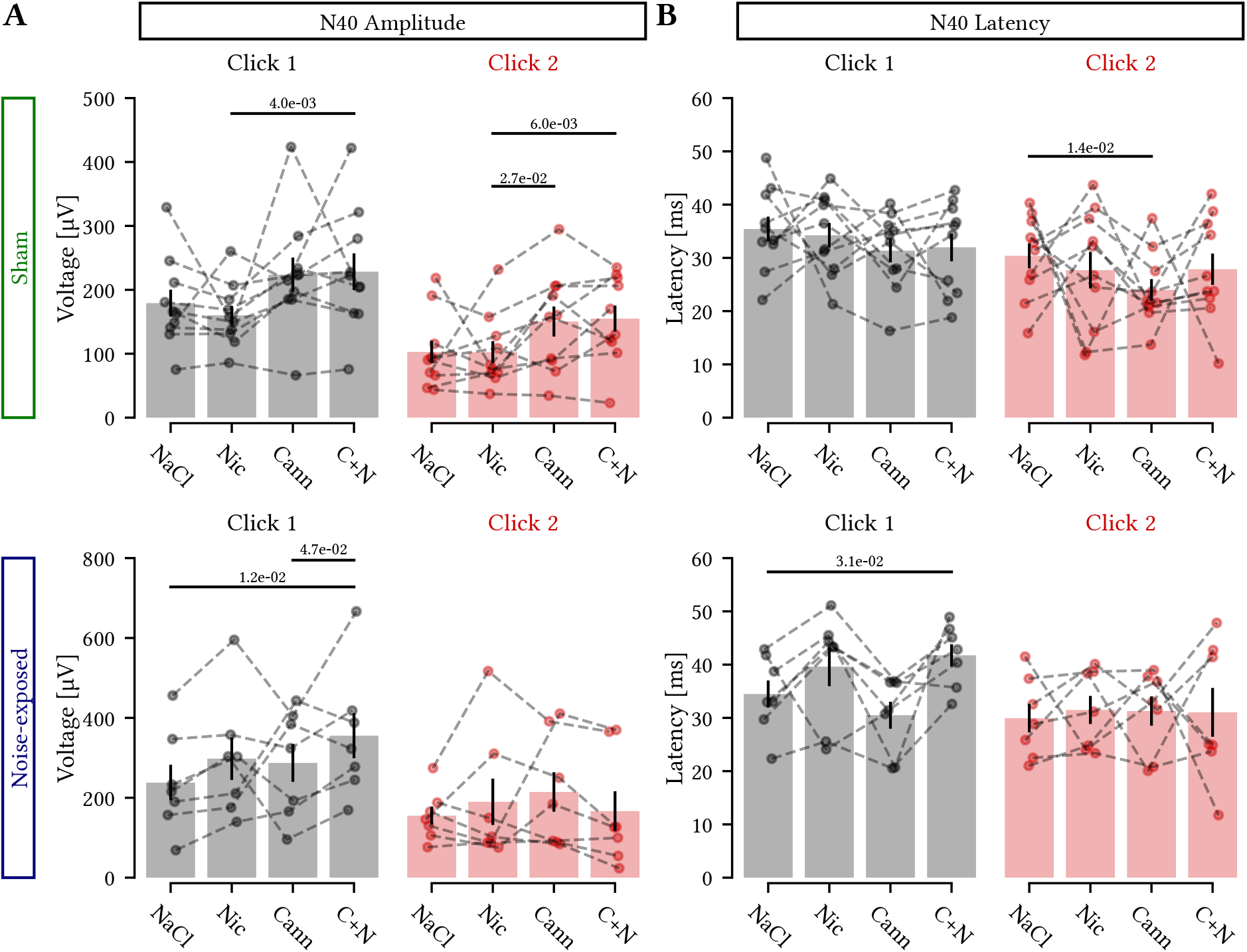
Noise-exposed mice only show modulation of the first click N40 response following cannabis+nicotine treatment. A) Comparison of the N40 amplitude in response to the first click (left) and second click (right) after saline, nicotine, cannabis extract and cannabis+nicotine administration for sham (top) and noise-exposed (bottom) mice. B) Latency comparisons between the first (left) and second (right) click responses in sham (top) and noise-exposed (bottom) animals across treatments. Only sham animals showed alterations of the second click amplitude and latency upon nicotine and cannabis treatment. Student’s t-test (A) and Wilcoxon signed-rank test (B), n = 10 sham and 7 noise-exposed mice.

Lastly we quantified the inter-peak interval (latency between the N40 and P80 peaks) of the response to the paired clicks (Supplemental Figure S5). When double peaks were present, we measured latency from the first peak in the doublet (Supplemental Figure S5A). We did not see any difference in the number of double N40 peaks recorded from sham and noise-exposed animals (p *>* 0.07 for all conditions tested; Supplemental Figure S5B). Also, there were no significant differences in the inter-peak interval between negative and positive aERP for either treatments or groups (F(1,20) *<* 2.06, p *>* 0.1; Supplemental Figure S5C). Thereby the average aERP waveform appears robust for latencies, despite individual variability.

Taken together, this study found noise-exposed mice to normally gate repetitive auditory stimuli, but showing larger amplitudes and slower processing of attention to repetitive clicks after pharmacological perturbations of the cholinergic and endocannabinoid systems, compared to sham-treated animals. The modulation of aERPs under cannabis + nicotine treatment was specifically related to the first click of the N40 component amplitude in noise-exposed mice.

## Discussion

We found that the N40 amplitude and latency is increased in animals with mild noise-exposure (Figure 7A-B). These mice showed increased ratio of the amplitude of first and second click N40 components upon cannabis and nicotine administration compared to sham animals (Figure 7C), which indicates improvement in sensory gating. Cannabis administration also increased the latency ratio of the N40 component of aERPs for noise-exposed mice compared to sham mice (Figure 7D), indicating altered temporal processing. Our findings imply that cholinergic and endocannabinoid signaling and/or downstream pathways are involved in perturbed sound processing after mild noise exposure. Still, the cannabis extract may contain substances that act on non-endocannabinoid targets [47], and further studies utilizing isolated endocannabinoid receptor agonists could elucidate the involvement of these receptors in sound processing.

**Figure 7:**
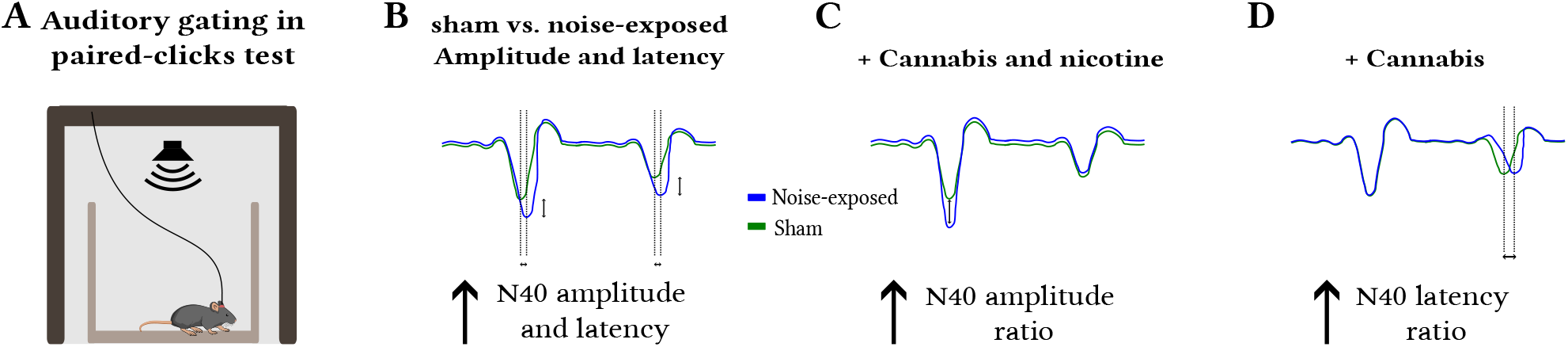
Schematics of the main findings. A) Experimental setup showing an implanted animal during the paired-click test recording. B) N40 amplitude and latency is increased in noise-exposed animals compared to sham. C) Cannabis + nicotine treatment improved N40 ratio by increasing the first click response. D) Cannabis treatment increased the second click latency ratio for noise-exposed animals compared to sham.

Tinnitus is a highly heterogeneous disorder in humans [48], and the underlying pathophysiological mechanisms remain unclear. Recent evidence in animals and humans cumulate towards the involvement of the limbic system in tinnitus [7], however the confounding effects of hearing loss and hyperacusis make the disentangling of each contributing factor quite challenging [49]. Our data is consistent with findings described by Campbell et al. (2018), studying young individuals with mild tinnitus and a normal audiogram. They found poorer auditory processing, indicating impaired sensory gating, due to no significant difference between response amplitudes of the first and second P50 aERP for tinnitus patients [19], similar to what we found for N40 under saline treatment (Supplemental Figure S2). Thereby our animal model results match patients with mild tinnitus. To our knowledge, this is the first study to investigate sensory gating in the hippocampus in noise-exposed mice and to evaluate how the cholinergic and endocannabinoid system interferes with sensory gating in these animals. A strength of this study is that hippocampal location for quantifying aERPs was standardized by anatomical post hoc examination and by electrophysiological profile [32] at each treatment session, thereby opening up for systematically testing a variety of compounds affecting limbic processing of attention to sound.

Another limitation is that the direct impact of nicotine and the cannabis extract on tinnitus were not assessed after the pharmacological intervention. This limitation was due to the size of the implanted electrode, thereby not allowing animals to enter the restraining tube, designed to make mice stand on all four paws during GPIAS measurements. Previous studies of cannabis as a tinnitus treatment have shown conflicting results [50, 51]. For instance, acute injection of the synthetic CB1/CB2 receptor agonists (WIN55,212-2, or CP55,940), exacerbate salicylate-induced tinnitus in rats assessed using a conditioned lick suppression paradigm [52], whereas acute treatment with the CB1 receptor agonist arachidonyl-2-chloroethylamide (ACEA) had no effect (as measured by GPIAS) in guinea pigs with salicylate-induced tinnitus [53]. It is possible that the confounding effects of stress on GPIAS measures caused by either salicylate or cannabis complexify the behavioral interpretation.

Based on the hypothesis that tinnitus can be similar to epilepsy due to hyperactivity in auditory and non-auditory pathways [54], here we used an extract containing a high dose of THC, since it was previously demonstrated that high THC doses presented anticonvulsant effects. 50mg/kg THC was shown to prevent spontaneous seizures in rodents [55, 56]; and THC doses up to 100 mg/kg, with effective dose at 48 mg/kg, to be anticonvulsant after seizure generation by electroshock in mice [57]. Even 80 mg/kg THC effectively suppressed pharmacologically-induced convulsions, delayed their onset, and prevented mortality in mice [58]. In addition, 96% of patients of a Canadian study reported that they would consider cannabis as treatment for their tinnitus [59]. Furthermore, cannabis extract concentration has shown U-shaped dose-response antidepressant effects in mice [40], thereby evaluating dose-dependent effects of activating CB1 receptors in different tinnitus models, as well as comparisons of administration routes of cannabis extract, is necessary in future studies.

Here we found that pharmacological manipulations of aERPs with both nicotine and cannabis extract improve sensory gating in noise-exposed mice but not in sham-treated animals. Our findings suggest that the higher N40 ratio under cannabis extract together with nicotine treatment in noise-exposed mice is related to an elevated click 1 amplitude and a lack of consistent modulation of the response to the second click, suggesting an increased registration (sensorial input processing) of the stimulus, as suggested previously [60]. Probably this effect was not seen in sham animals because both clicks were modulated by the treatments containing cannabis.

Nicotine is known to increase the amplitude of the P20 and N40 1st click in mice [13, 61]. The second click response has instead been shown to be sensitive to muscarinic receptor antagonists, increasing the second click amplitude and disrupting sensory gating [62]. Next, the P80 response is known to be reduced by NMDA receptor antagonists such as ketamine [46, 63]. Thereby, an active cholinergic system appears to facilitate auditory gating of the N40 response, but it is important to notice that smoking is associated with greater risk of tinnitus [3]. We speculate that for tinnitus models nicotine might suppress hyperactivity in the dorsal cochlear nucleus since it has been previously demonstrated that cholinergic agonists such as carbachol can suppress noise-induced hyperactivity in the DCN in rodents [64], possibly affecting sound processing in higher areas.

The combination of cannabis extract and nicotine could potentially cause interaction effects, since it has been shown in isolated cells that anandamide (an endogenous CB1 receptor agonist) decreased nicotinic currents generated by nicotinic *α*7 and *α*4*β*2 subunit containing acetylcholine receptors [65]. Also, a link between cannabis dependency and activity of subtypes of nicotinic acetylcholine receptors has recently been shown [66, 67]. Furthermore, the interplay between the cholinergic and endocannabinoid system has been shown in basal forebrain cholinergic neurons expressing CB1 receptors [68] and interestingly, human subjects administered orally a combination of a THC analog and nicotine have shown improved auditory deviant detection and mismatch negativity aERPs, but not when each drug was delivered alone [22]. Since we found only noise-exposed animals to improve N40 amplitude gating ratio in response to cannabis+nicotine treatment, and since it has been demonstrated that vesicular acetylcholine transporters puncta density is decreased on both sides of the hippocampus after noise exposure [69], we hypothesize that nicotine administration could be compensating for a decrease in acetylcholine release in these animals. Still, the cellular mechanisms underlying such alterations in sensory gating remain to be further investigated.

In general, the endocannabinoid system dampens neuronal activity by activation of Gi-protein coupled presynaptic CB1 receptors that decrease neurotransmitter release through blocking of presynaptic voltage-gated calcium channels and opening of voltage-gated potassium (GIRK) channels, allowing potassium to flow out of the terminal [70]. For example, high doses of natural cannabis extracts can reduce neuronal hyperactivity in in vitro models of spasticity and epilepsy [71] which is interesting since noise-induced tinnitus is related to neuronal hyperactivity of the auditory system [9]. Still, the circuit effect of CB1 receptor activation depends on what type of presynaptic neuron expresses CB1 receptors (etc. glutamatergic or GABAergic cells) which can affect local plasticity differently [72]. It is known that pyramidal cells of the hippocampus have relatively low expression of CB1 receptors [73] therefore we expect the cannabis extract to increase auditory input due to decreased inhibition, since CB1 receptors are strongly coexpressed with GAD65 in the hippocampus [73, 74], especially with strong CB1 receptor expression on cholecystokinin positive interneurons [74]. Furthermore, this study uses a THC-rich extract, which needs to be put in contrast to anxiolytic evaluation of THC at much lower doses [39] and studies of seizure reduction by THC at doses as high as 100 mg/kg [55]. Still the concentration of THC in a cannabis extract cannot be compared to THC alone, but should be considered in relation to other cannabinoids present. For example, a systematic review of cannabinoid treatment of chronic pain found products with high-THC-to-CBD ratios the most useful for short-term relief of neuropathic chronic pain [75].

The ability to suppress repetitive auditory stimuli was preserved in noise-exposed mice, suggesting that noise-induced tinnitus without changes in hearing thresholds does not interfere with auditory gating but that noise-induced tinnitus renders the response to auditory clicks abnormal in the presence of cannabis by delaying temporal coding. Here we found that cannabis alone did not decrease aERP amplitude as has seen in human P300, probably due to the N40 component (human N100) reflecting triggered attention [76] and the human P300 reflecting cognitive stimulus classification [21]. It is important to pin-point cellular contribution to the aERP components and here, due to the availability of a transgenic line targeting Cre expression at cells expressing the alpha-2 nicotinic receptor [ChRNA2; 77], the role of the cholinergic system in sensory gating and tinnitus could be investigated by using chemogenetics to locally manipulate the excitability of these cells during aERP recordings; or in tinnitus induction performing similar manipulations during noise exposure. A similar approach would be difficult for investigating the role of the endocannabinoid system in tinnitus due to the unavailability of specific targeting of, for example, CB1-expressing cells. However, the depletion of glutamate aspartate transporter (GLAST) to exacerbate the tinnitus phenotype, may also be more appropriate to investigate in greater details the underlying cellular and molecular mechanisms [78]. Still, it is becoming clear that loud noise activates both auditory and limbic pathways [79] but how prolonged noise-exposure alters sound processing of each system needs to be further examined, as well as how the limbic and auditory systems interact in tinnitus [11].

In conclusion, our study shows that provoking auditory event-related potentials pharmacologically, using nicotine and/or cannabis extract rich in THC, showed noise-exposed mice to improve gating of the N40 component especially under the combined influence of cannabis extract and nicotine, by increasing the first click response amplitude. However, cannabis extract also increased the latency ratio of the N40 component in noise-exposed mice compared to sham animals, indicating delayed temporal processing of paired clicks. Thereby the activation of the cholinergic and endocannabinoid receptors and downstream pathways have distinct and different effects on auditory gating in the context of tinnitus phenotype. Our findings provide insights into the neural processing alterations associated with tinnitus-like behavior, which may facilitate the future development of diagnostic methods and potential pharmacological interventions.

## Author Contributions

KEL and BC designed the study. BC and TM performed experiments; SRBS analyzed the cannabis extract; BC, TM and TZL analyzed the data; BC, CRC and KEL wrote the manuscript with important input from TM and TZL.

## Acknowledgments

The authors thank Dr Claudio Queiroz at the Brain Institute-UFRN for providing the cannabis extract and Dr Jessica Winne for technical assistance in electrode manufacturing.

## Data Availability Statement

The datasets generated and/or analyzed in the current study are available on request. Recordings were done using the Open-ephys GUI [80]. Stimulation and data analysis were performed using SciScripts [81], Scipy [82] and Numpy [83]. All plots were produced using Matplotlib [84], and schematics were done using Inkscape [85]. All scripts used for recordings and analysis are available online [86].

