## Supplemental File for "Alterations of auditory sensory gating in mice with noise-induced tinnitus treated with nicotine and cannabis extract"

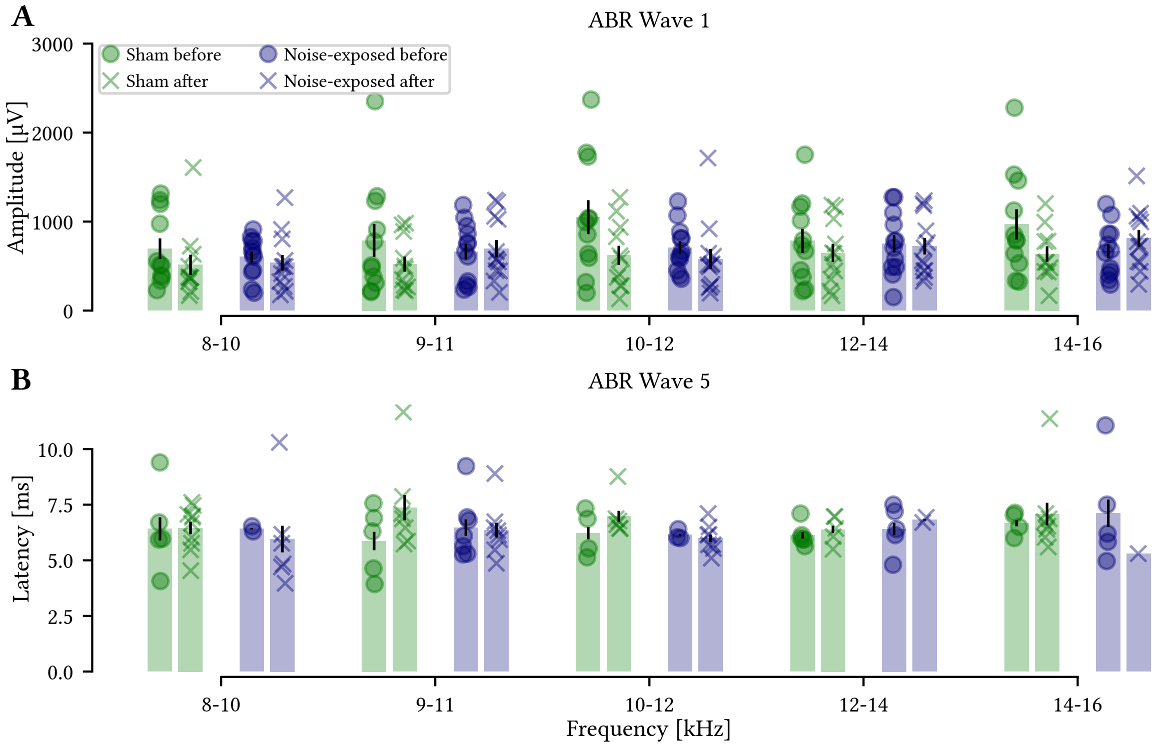


**Supplemental Figure S1.** ABRs showed no alteration of wave 1 amplitude or wave 5 latency in sham or noise-exposed mice. Wave 1 amplitude (A) and wave 5 latency (B) for each frequency showing no difference before and after noise-exposure for both sham (green) and noise-exposed (blue) mice. Wilcoxon signed-rank test, n = 11 sham and 11 noise-exposed mice.

**Supplemental Table S1**. 3-way ANOVA evaluating the effect of the factors Group (sham vs noise-exposed), Treatment (saline, nicotine, cannabis, cannabis+nicotine) and Click repetition (1st or 2nd click) for N40 amplitude.

| Effect | DFn | DFd | F | p | p<.05 |
|---|---|---|---|---|---|
| Group | 1 | 20 | 7.467 | 6.3e-03 | * |
| Treatment | 3 | 60 | 3.245 | 2.8e-02 | * |
| Click | 1 | 20 | 21.903 | 1.4e-04 | * |
| Group:Treatment | 3 | 60 | 0.875 | 0.459 | |
| Group:Click | 1 | 20 | 3.194 | 0.089 | |
| Treatment:Click | 3 | 60 | 3.517 | 2.0e-02 | * |
| Group:Treatment:Click | 3 | 60 | 3.668 | 1.7e-02 | * |

**Supplemental Table S2**. Statistics evaluating the effect of the factors Group (sham vs noise-exposed), Treatment (saline, nicotine, cannabis, cannabis+nicotine) and Click repetition (1st or 2nd click) on N40 latency.

| Method | Effect | DFn | DFd | eff. size | p | p<.05 |
|---|---|---|---|---|---|---|
| Kruskal-Wallis | Group | 1 | 20 | 4.11 | 4.3e-02 | * |
| Friedman | Treatment | 3 | 60 | 0.858 | 0.468 | |
| Friedman | Click | 1 | 20 | 7.87 | 2.6e-03 | * |

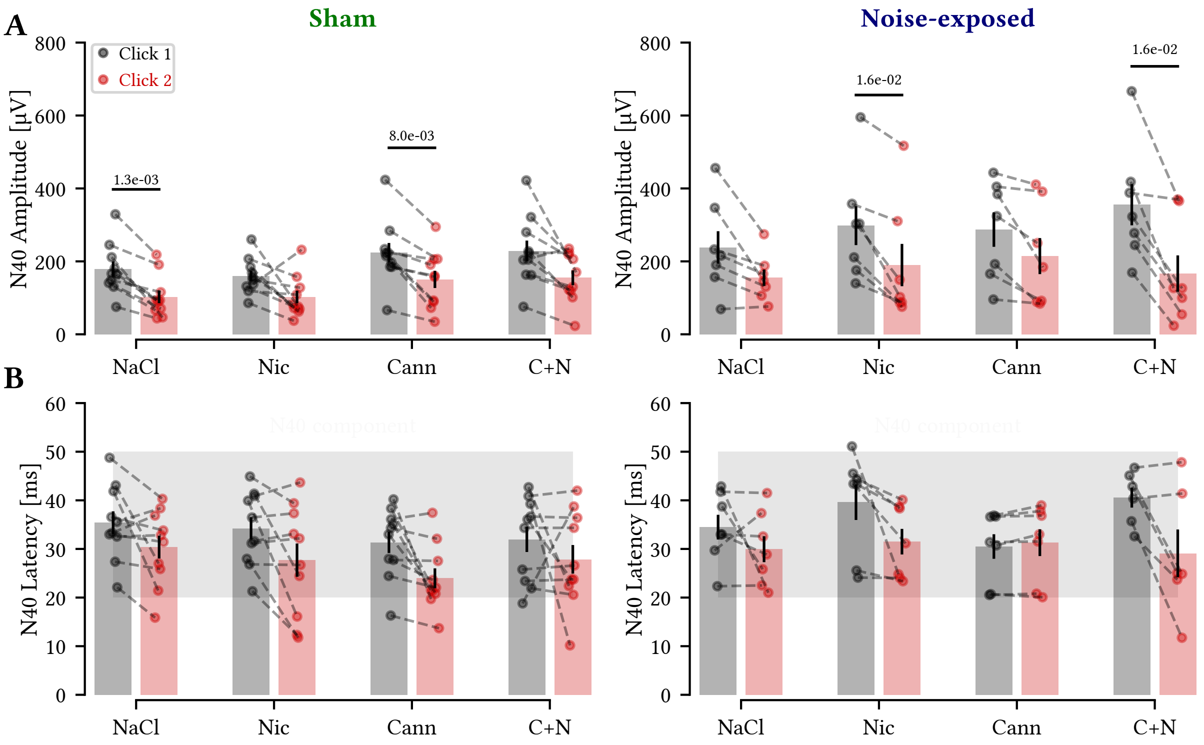


**Supplemental Figure S2.** Noise-exposed animals showed decrease in average second click amplitudes compared to the first click under nicotine and cannabis extract + nicotine treatment. A) Quantification of the N40 amplitude in response to the first (gray) and second (red) clicks in different pharmacological treatments for sham (left) and noise-exposed (right) mice. Sham-exposed mice displayed a decrease in response to the second click in the presence of saline ($p = 1.3e\text{-}03$) and nicotine removed the difference between the average first and second click, while in the presence of cannabis extract a difference in click 1 and and 2 amplitudes was maintained ($p = 8.0e\text{-}03$) but gone if cannabis extract and nicotine treatment was combined. For noise-exposed mice responses to the first and second click were not significantly different in NaCl treatment ($p = 0.237$), while significant different in the presence of nicotine ($p = 1.6e\text{-}02$) and after cannabis extract + nicotine treatment ($p = 1.6e\text{-}02$). B) The latency of the first and second click was not affected by different treatments in neither sham nor noise-exposed mice. The negative peak (N40) occurred within 20-50ms latency (gray shading), with a smaller latency for the second peak in most cases. Student's t-test (A) and Wilcoxon signed-rank (B); n = 10 sham and 7 noise-exposed mice.

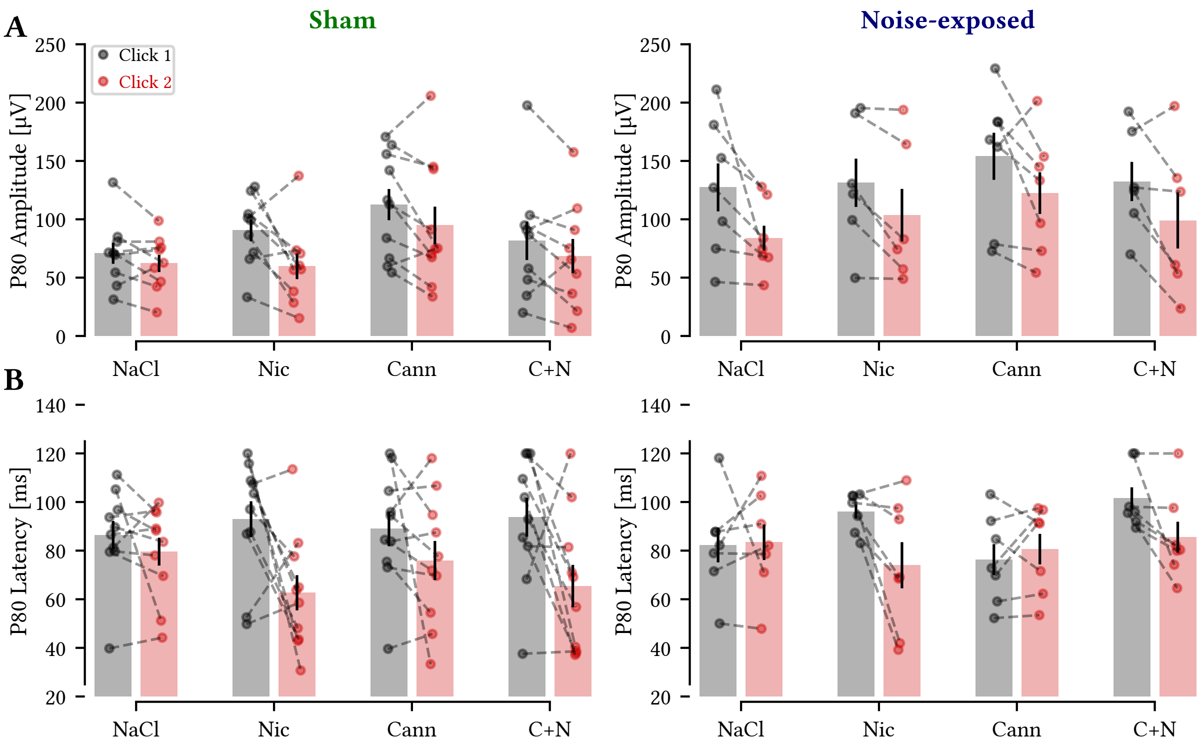


**Supplemental Figure S3.** The P80 component was not altered by noise or pharmacological treatment. A) The aERP component P80 amplitude was not altered by nicotine and/or cannabis extract for sham (left) or noise-exposed (right) mice. B) There was a large variability in latency of the positive peak for both sham and noise-exposed animals, with no significant differences between groups at any treatment. Student's t-test, n = 10 sham and 7 noise-exposed mice.

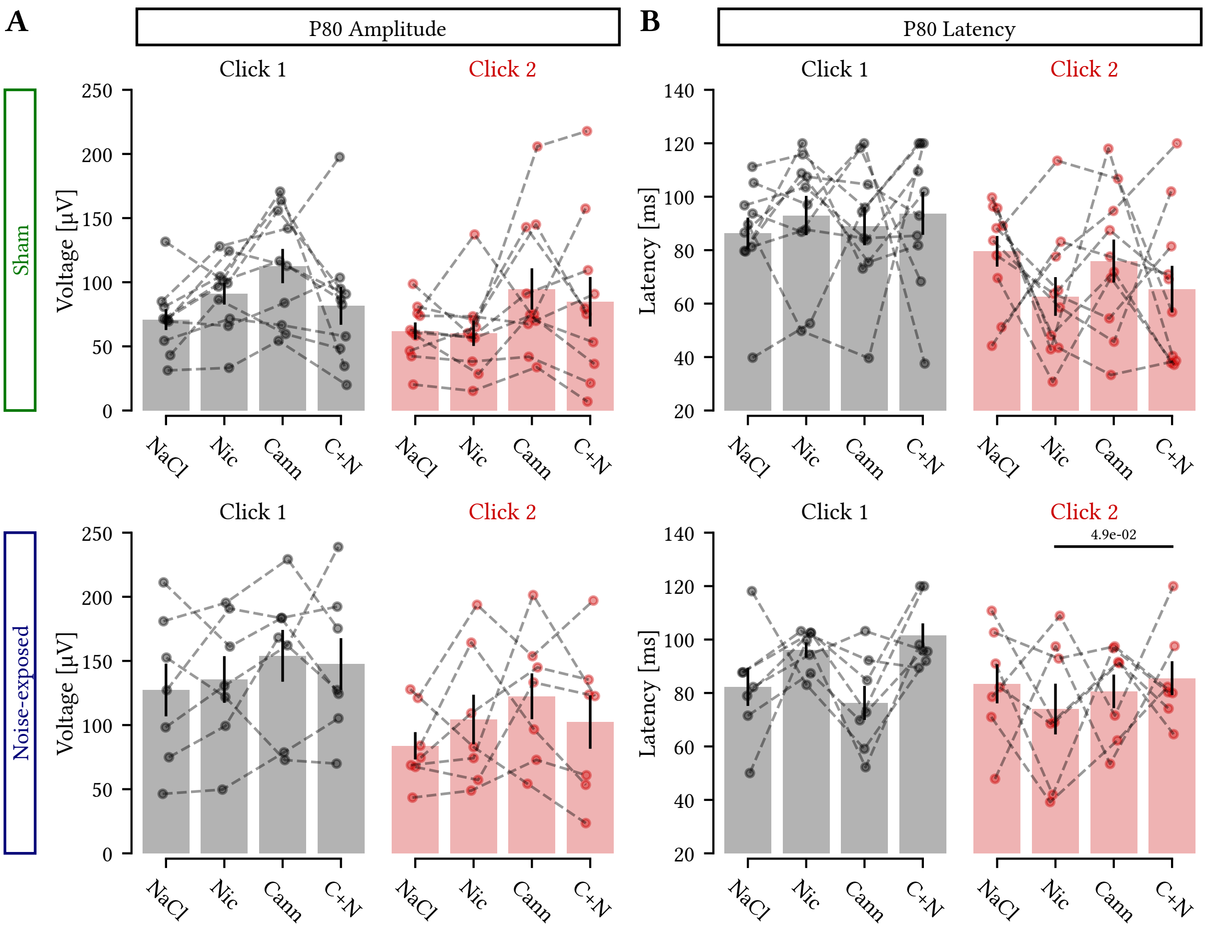


**Supplemental Figure S4.** Pharmacological treatment did not affect the P80 first or second click amplitude. A) Click 1 (top) and click 2 (bottom) P80 amplitude response for sham and noise-exposed tinnitus mice under different pharmacological treatments. B) Same as for 'A' for P80 latency. Only the combination of cannabis and nicotine was marginally delaying the second click compared to only nicotine for noise-exposed mice. Student's t-test, n = 10 sham and 7 noise-exposed mice.

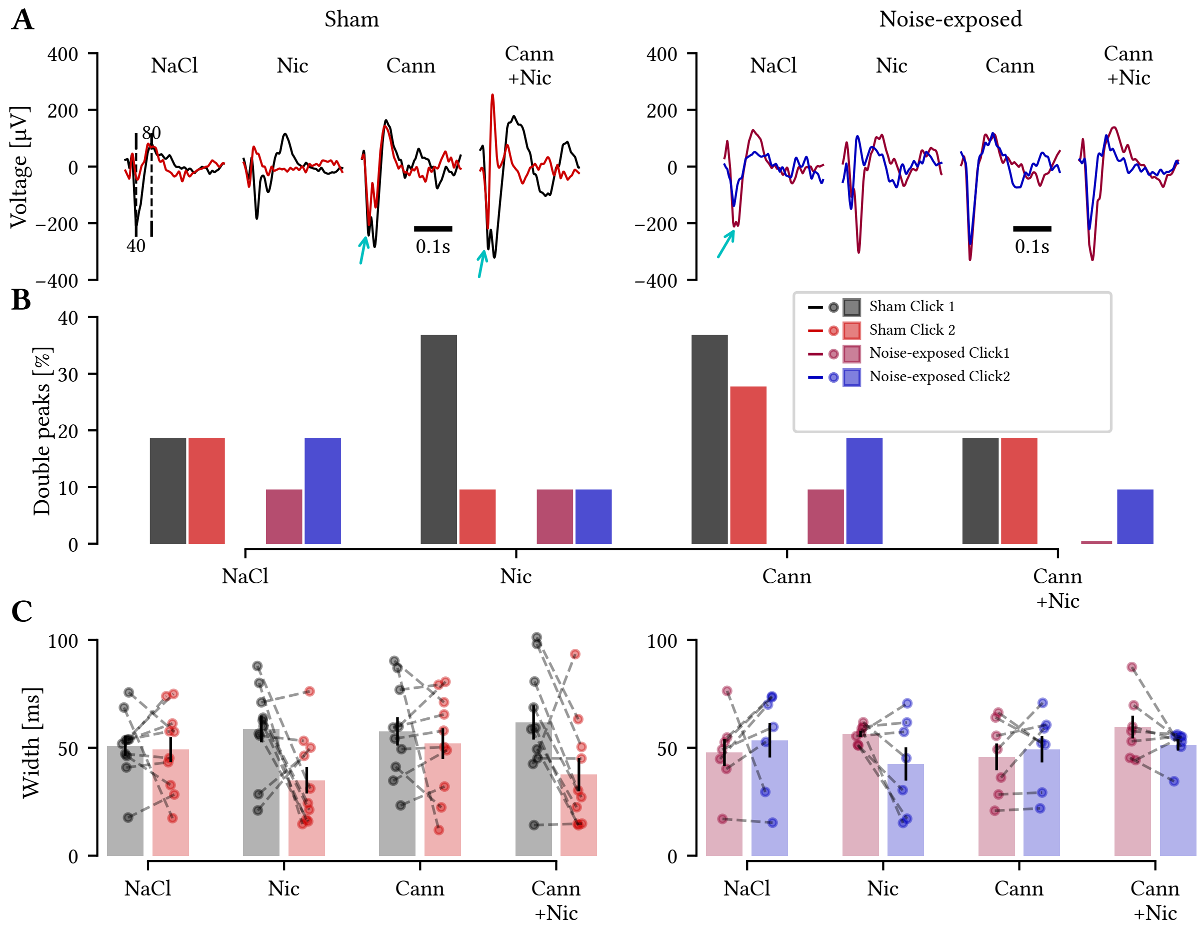


**Supplemental Figure S5.** Noise exposure and pharmacological treatments do not affect aERP interpeak interval. A) Representative aERP traces for a sham (left) and a noise-exposed animal (right) of the first (black and purple) and second (red and blue) click with occasional doublet peaks (see cyan arrows). B) Percentage of double N40 peaks in sham and noise-exposed mice showed no difference in prevalence ($p > 0.07$ for all conditions tested, McNemar's test). C) Inter-peak interval (time between N40 and P80 components) for the first (black and purple) and second (red and blue) click response for sham (left) and noise-exposed (right) animals shows that the time between negative and positive peaks varies but the average is closer to 50ms rather than 40ms as indicated by the literature nomenclature. Student's t test, $n = 10$ sham and 7 noise-exposed mice; $p > 0.05$ for all comparisons.
